# High-performance GFP-based calcium indicators for imaging activity in neuronal populations and microcompartments

**DOI:** 10.1101/434589

**Authors:** Hod Dana, Yi Sun, Boaz Mohar, Brad Hulse, Jeremy P. Hasseman, Getahun Tsegaye, Arthur Tsang, Allan Wong, Ronak Patel, John J. Macklin, Yang Chen, Arthur Konnerth, Vivek Jayaraman, Loren L. Looger, Eric R. Schreiter, Karel Svoboda, Douglas S. Kim

## Abstract

Calcium imaging with genetically encoded calcium indicators (GECIs) is routinely used to measure neural activity in intact nervous systems. GECIs are frequently used in one of two different modes: to track activity in large populations of neuronal cell bodies, or to follow dynamics in subcellular compartments such as axons, dendrites and individual synaptic compartments. Despite major advances, calcium imaging is still limited by the biophysical properties of existing GECIs, including affinity, signal-to-noise ratio, rise and decay kinetics, and dynamic range. Using structure-guided mutagenesis and neuron-based screening, we optimized the green fluorescent protein-based GECI GCaMP6 for different modes of *in vivo* imaging. The jGCaMP7 sensors provide improved detection of individual spikes (jGCaMP7s,f), imaging in neurites and neuropil (jGCaMP7b), and tracking large populations of neurons using 2-photon (jGCaMP7s,f) or wide-field (jGCaMP7c) imaging.

## Introduction

Measurement of neuronal activity using genetically encoded calcium indicators (GECIs) has become a widely used method in neuroscience, driven by concomitant improvements in both GECI performance and microscopy methods. For example, the GFP-based GCaMP sensors^1–3^ have been iteratively engineered to enhance the signal-to-noise ratio (SNR) for detection of neural activity. The widely used GCaMP6 sensors enable the detection of single action potential (AP) firing under favorable conditions^4^. They can be used to monitor the activity of large groups of neurons *in vivo* using two-photon microscopy^5,6^ and wide-field imaging^6^. They can also measure activity-induced calcium changes in small subcellular compartments like dendritic spines^4^ and axons^7^. GECIs allow measurement of neuronal activity over timescales of tens of milliseconds^8–10^ to months^11,12^.

Different types of imaging benefit from GECIs with specific properties. Imaging from large populations of neurons in densely labeled samples is helped by lower baseline fluorescence, which reduces the background signal from neuropil and inactive neurons. Low baseline fluorescence is especially important for wide-field fluorescence methods, where much larger tissue volumes contribute to the background. Neuronal microcompartments such as spines and axons present different challenges. Their small diameters (~0.1 μm) mean that few fluorescent molecules are present and molecules are more vulnerable to photobleaching. In addition, image quality is compromised by motion artifacts. Taken together, it can be hard to collect sufficient signal even to localize such compartments above the dark background. A sensor with substantial baseline fluorescence is therefore preferred for this application. High baseline fluorescence can also be helpful for imaging in sparsely labeled samples, where background signal is negligible, such as those produced by highly selective genetic drivers in *Drosophila*^13^. In addition, different applications demand sensors with different dynamics. Slow sensors are better for detecting activity, whereas fast sensors track the dynamics of physiological signals more faithfully.

We present a new generation of GCaMP sensors tailored to detect neuronal activity in specific circumstances. jGCaMP7 indicators include: jGCaMP7s (sensitive and slow), jGCaMP7f (fast kinetics), jGCaMP7b (brighter baseline fluorescence), and jGCaMP7c (high contrast with low baseline fluorescence). The jGCaMP7 sensors were tested *in vitro* and *in vivo*, and show substantially better performance than the GCaMP6 sensors.

## Results

### Sensor engineering

Crystal structures of GCaMP in the presence or absence of Ca^2+^ ^14,15^ were used to specify positions likely to affect key sensor properties^2–4^, including resting and peak fluorescence, and affinity and kinetics of Ca^2+^ binding and unbinding^16^. GCaMP comprises a circularly permuted GFP (cpGFP) fused to both calmodulin (CaM) and a calmodulin-binding peptide from myosin light-chain kinase (RS20, also known as M13)^1^. Calcium-dependent conformational changes in CaM and its interactions with both M13 and GFP itself modulate the GFP chromophore environment and affect fluorescence^17,18^. We targeted mutagenesis to the GFP-CaM interface and the CaM-M13 peptide interaction (Fig. 1)^3,4^. In all, 27/450 amino acid positions (mostly sites not mutated in previous cycles) were mutated to near-saturation (Supp. Table 1). In addition, the M13-to-cpGFP inter-domain linker, that increased sensitivity in the GCaMP5 variant series^3^, was reintroduced, and the cpGFP-to-CaM linker was altered. Mutations were made on top of GCaMP5G, GCaMP6s, or GCaMP6f.

**Figure 1.**
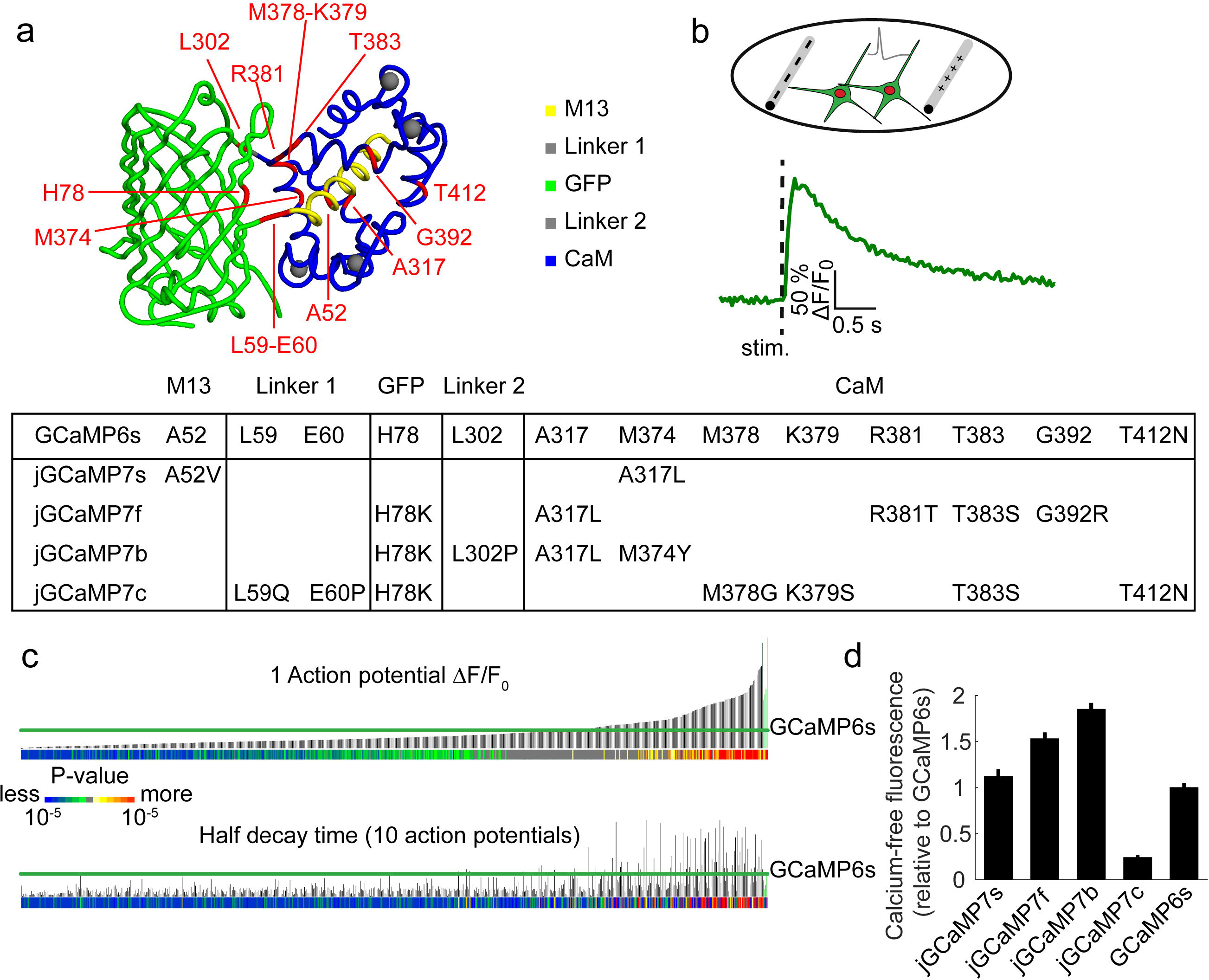
Mutagenesis and screening of jGCaMP7 in dissociated neurons. **a**, Mutations introduced in jGCaMP7 sensors (top). M13 peptide (yellow), linker 1 and 2 (gray), cpGFP (green) and CaM (blue). Table, mutations for each jGCaMP7 variant. **b**, Top, schematic cultured neurons with stimulation electrodes (gray). Cultured neurons expressed a cytosolic jGCaMP7 variant and nuclear mCherry. Bottom, change in fluorescence of a single well with cells expressing jGCaMP7s after firing one action potential (AP). **c**, Top, screening results for 662 jGCaMP7 variants. Top, fluorescence changes in response to one action potential (vertical black bars, ΔF/F_0_ amplitudes of tested variants; green bar, ΔF/F_0_ amplitudes of jGCaMP7c, f, b, and s, from left to right respectively). Horizontal green lines indicates GCaMP6 performance levels. Colored bars show the p-value of increase (hot colors) or decrease (cold colors) compared to GCaMP6s. Bottom, half decay times after 10 APs. **d**, Baseline fluorescence (mean±s.d) for jGCaMP7 and GCaMP6 sensors for purified proteins without calcium.

### Screening and *in vitro* characterization of jGCaMP7

Sensor variants (662 in all) were tested in cultured neurons ^19,20^. Neonatal rat hippocampi were dissociated, transfected with DNA constructs, and cultured in 96-well plates for 16-18 days. Trains of action potentials (APs) were triggered by electrical field stimulation within each well (Fig. 1b, Methods). Time-lapse fluorescence images (800 μm × 800 μm fields of view; 35 Hz) were acquired before, during, and after stimulation. Calcium-dependent fluorescence changes (ΔF/F_0_) were extracted from single neurons. The resting fluorescence (F_0_), sensitivity (*i.e.* fluorescence response to small stimuli), dynamic range (*i.e.* the response range between 1 AP and 160 AP), and kinetics were quantified for each variant (Supp. Table 1). Previously characterized GCaMP3, GCaMP5G, GCaMP6s, GCaMP6f, G-GECO1.0, and G-GECO1.1 were included in the screening as standards.

**Table 1.**
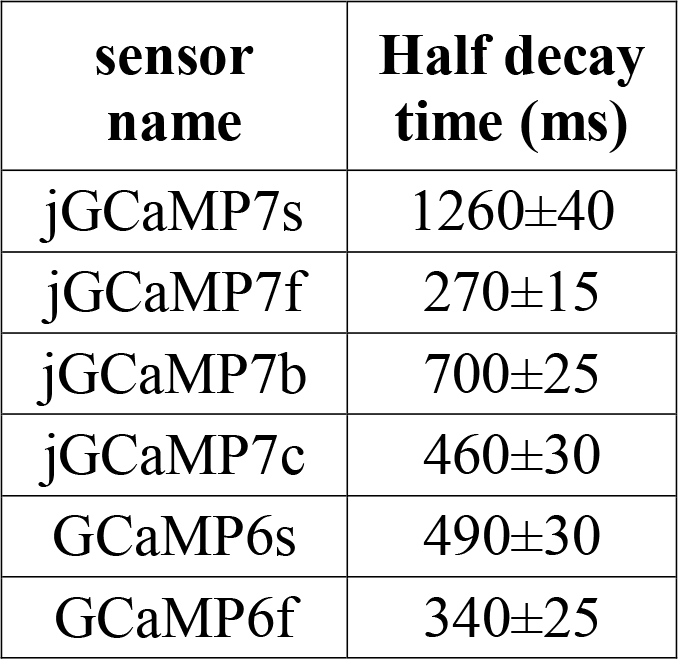
Half decay times of fluorescence signal *in vivo*

Many single mutations improved sensitivity compared to the parent proteins: 77/662 sensors showed higher (P < 0.01, Wilcoxon rank-sum test) 1-AP ΔF/F_0_ than GCaMP6s (Fig. 1c-top). About half of these sensitive variants exhibited slower decay times; other variants were simultaneously more sensitive and faster than GCaMP6s (Fig. 1c-bottom). Variants were sorted based on improved ΔF/F_0_ (in response to trains of 1, 3, 10, and 160 APs) and/or faster kinetics. Beneficial mutations were combined for a second round of screening.

Four new jGCaMP7 (“Janelia GCaMP7”, not to be confused with G-CaMP7^21^) sensors with distinct properties were selected (Fig. 2): 1) jGCaMP7s (“sensitive”) produces the highest response to small (1-10) AP trains. jGCaMP7s has a 5-fold larger ΔF/F_0_ amplitude for 1 AP stimuli and faster rise time than GCaMP6s. 2) jGCaMP7b shows a 3-fold increase in 1 AP ΔF/F_0_ response and a 50% increase in resting fluorescence compared to GCaMP6s. 3) jGCaMP7c (“contrast”) exhibits lower resting fluorescence, 2.7-fold increase in 1AP response, and greater ΔF/F_0_ to longer (20-160) AP trains compared to GCaMP6s, 4) jGCaMP7f (“fast”) has 5- and 3- fold larger 1-AP ΔF/F_0_ than GCaMP6f and GCaMP6s, respectively, with intermediate kinetics (1-AP half rise time: GCaMP6f, 26±2 ms (median±s.e.m.); jGCaMP7f, 26±2 ; GCaMP6s, 58±3. 1-AP half decay time: GCaMP6f, 140±20 ms; jGCaMP7f, 265±20; GCaMP6s, 455±40) (Supp. Table 2).

**Figure 2.**
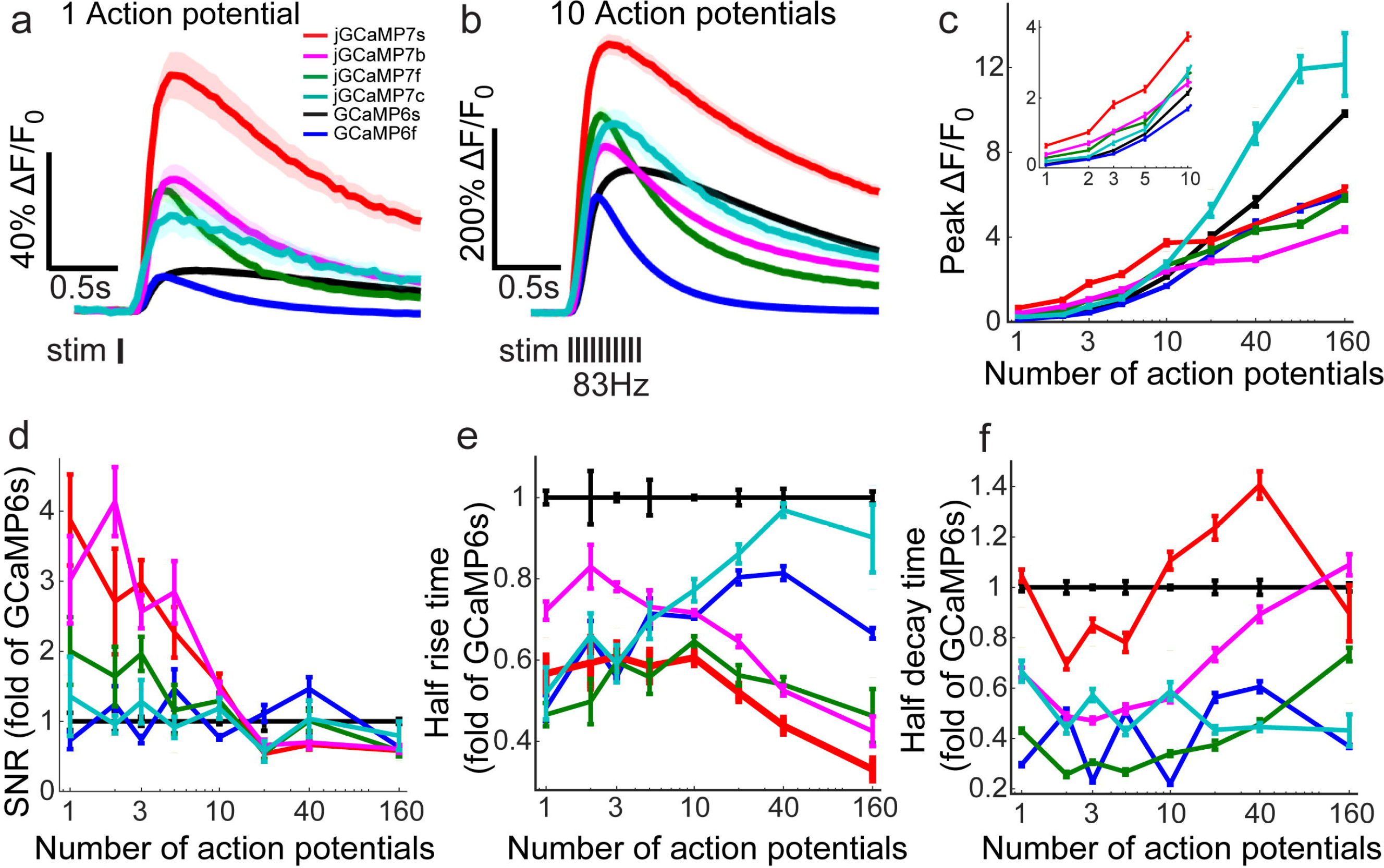
jGCaMP7 performance in dissociated neurons. **a**, Average responses to one action potential for jGCaMP7s (2173 neurons, 76 wells), jGCaMP7b (2986 neurons, 104 wells), jGCaMP7f (2369 neurons, 75 wells), jGCaMP7c 1379 neurons, 54 wells), GCaMP6s (17398 neurons, 682 wells), and GCaMP6f (19181 neurons, 673 wells). **b**, Same for 10 APs. **c-f**, Comparison of jGCaMP7 and GCaMP6 sensors, as a function of number of APs (color code as in **a**). **c**, Response amplitude, ΔF/F_0_. **d**, Signal-to-noise ratio, SNR, defined as the peak fluorescence, divided by the signal standard deviation before the stimulus. **e**, Half rise time. **f**, Half decay time. Error bars correspond to s.e.m (for 1,3, 10, and 160 field potentials, n=76 wells for jGCaMP7s; 104, jGCaMP7b; 75, jGCaMP7f; 54, jGCaMP7c; 682, GCaMP6s; 673, GCaMP6f. For 2,5,10,20, and 40 field potentials, n=27, jGCaMP7s;22, jGCaMP7b; 29, jGCaMP7f; 27, jGCaMP7c; 41, GCaMP6s; 41, GCaMP6f) (see Methods, see Supp. Table 4 for full datasets).

The GECI variants were purified for detailed *in vitro* spectroscopic and kinetic characterization. Consistent with its greater sensitivity in neurons, jGCaMP7s showed a lower dissociation constant (higher affinity) for Ca^2+^ binding compared to GCaMP6s (68 ± 3 nM vs. 147 ± 5 nM for GCaMP6s), mediated largely by a larger on-rate (k_on_; 21.5 ± 0.95 M^−1^s^−1^ vs. 4.3 ± 0.2 M^−1^s^−1^) (Supp. Table 1). Purified Ca^2+^-free jGCaMP7b was almost twice as bright (mediated by a higher extinction coefficient, ε_apo_) as GCaMP6s, with higher affinity, consistent with the neuronal data. Purified Ca^2+^-free jGCaMP7c was 25% as bright as GCaMP6s, mediated by a lower ε_apo_ and a significantly increased pK_a_-apo (8.66 ± 0.05 vs. 7.54 ± 0.08). Purified jGaMP7f showed fast k_on_ and k_off_ for Ca^2+^ binding (Supp. Table 1). For all variants, one- and two-photon fluorescence absorption and emission spectra are similar to the parent constructs (Supp. Figs. 1-2).

### Imaging in the *Drosophila* larval neuromuscular junction

We imaged jGCaMP7 sensors in presynaptic boutons of the *Drosophila* larval neuromuscular junction (Fig. 3a). jGCaMP7 sensors showed improved sensitivity and speed compared to GCaMP6. jGCaMP7s exhibited 2.5 to 5-fold higher ΔF/F_0_ response amplitudes compared to GCaMP6s at low frequency stimulation (1-10 Hz; Fig. 3b-d, g-h; Supplementary Table 3), but jGCaMP7s responses were lower than those of GCaMP6s with high frequency stimulation (40- 160 Hz; Fig. 3f-h; Supplementary Table 3). jGCaMP7s was generally slower than GCaMP6s (Fig. 3j-k). jGCaMP7f displayed similar response amplitudes compared to GCaMP6f in the low frequency range (1-10 Hz; Fig. 3b, d, g-h; Supplementary Table 3) but was 25 to 50% faster in terms of decay kinetics (Fig. 3j; Supplementary Table 3). jGCaMP7b exhibited 2 to 4-fold higher amplitude at low frequencies and 50% higher resting fluorescence compared to GCaMP6s (Fig. 3b, d, g-h, l). jGCaMP7c exhibited higher ΔF/F_0_ responses than GCaMP6s with both low and high frequency stimulation (Fig. 3b, d, f-h; Supplementary Table 3). Its resting fluorescence was about one third of the GCaMP6s level (Fig. 3l), but the signal-to-noise ratio of the two sensors was similar (Fig. 3i).

**Figure 3.**
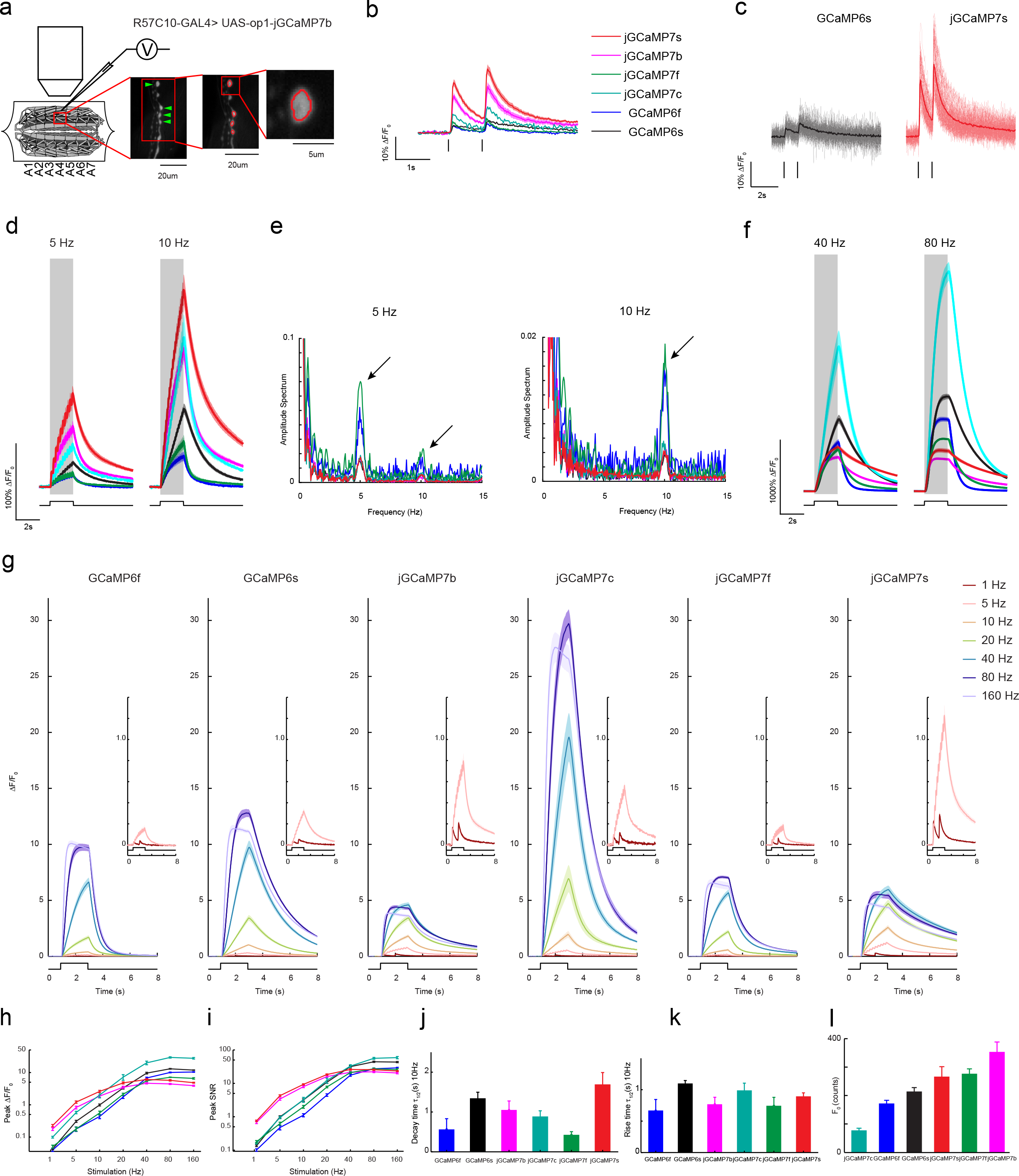
jGCaMP7 performance in the *Drosophila* larval neuromuscular junction. (**a**) Experimental setup. Epifluorescence images of Type 1b boutons (green arrowheads) from muscle 13 (segments A3-A5), with image segmentation ROIs superimposed (Materials and methods). (**b**) Averaged fluorescence transients responding to two individual APs (ticks at the bottom) (jGCaMP7s: 14 FOVs in 7 flies, 56 boutons; jGCaMP7b: 12 FOVs in 7 flies, 48 boutons; jGCaMP7f: 14 FOVs in 7 flies, 56 boutons; jGCaMP7c: 12 FOVs in 7 flies, 48 boutons; GCaMP6f: 13 FOVs in 7 flies, 52 boutons; GCaMP6s: 14 FOVs in 7 flies, 56 boutons. Same data set used for all other analyses). Lines, mean; shading, s.e.m. (**c**) Single trial (thin lines) and mean (thick lines) responses to two APs for GCaMP6s and jGCaMP7s. (**d**) Comparison of averaged responses to 2s of 5 Hz and 10 Hz stimulation. Color code as in b. (**e**) Power spectra normalized to values at 0 Hz for fluorescence signals acquired during 5 Hz and 10 Hz stimulation. Color code as in b. (**f**) Comparison of averaged responses to 2s of 40Hz and 80Hz stimulation. Color code as in b. (**g**) Responses to 1, 5, 10, 20, 40, 80 and 160 Hz stimulation for 2 s. Insets, 1 and 5 Hz stimulation for 2 s. Lines, mean; shading, s.e.m. (**h–i**) Averaged peak △F/F_0_ (**k**) and peak SNR (**l**) as a function of frequency. Vertical axes are log scale. Error bars represent s.e.m. in h-l. (**j-k**) Comparison of decay (**j**) and rise (**k**) kinetics with 10 Hz stimulation. (**l**) Comparison of baseline fluorescence.

### Imaging a population of compass neurons in adult *Drosophila* during behavior

Imaging calcium activity from small, sparsely-labeled samples is often limited by the fluorescence signal that can be collected. This favors sensors with both a high baseline fluorescence, which aids in localization and reducing motion artifacts, and the ability to generate a large number of signal photons above baseline, which allows for lower excitation light intensity at equivalent signal-to-noise ratios. To assess how the jGCaMP7 variants perform under such circumstances, we imaged a population of neurons, known as E-PGs^22^, whose processes innervate a small (~50 μm diameter), doughnut-shaped neuropil called the ellipsoid body (Fig. 4b). During imaging, head-fixed flies walked on a spherical treadmill in closed loop with their visual surroundings (Fig. 4a). Previous work^10^ has shown that activity in this circuit is organized as a single ‘bump’ whose angular position encodes the fly’s heading direction (Fig. 4c-d). GCaMP6f, jGCaMP7f, jGCaMP7b, and jGCaMP7s all track this activity bump during virtual navigation, but with important differences (Fig. 4d). jGCaMP7f, jGCaMP7b and jCaMP7s have a higher baseline fluorescence and also generate larger signal changes above baseline (ΔF) compared to GCaMP6f (Fig. 4e-f), making them better suited for imaging small, dim structures.

**Figure 4.**
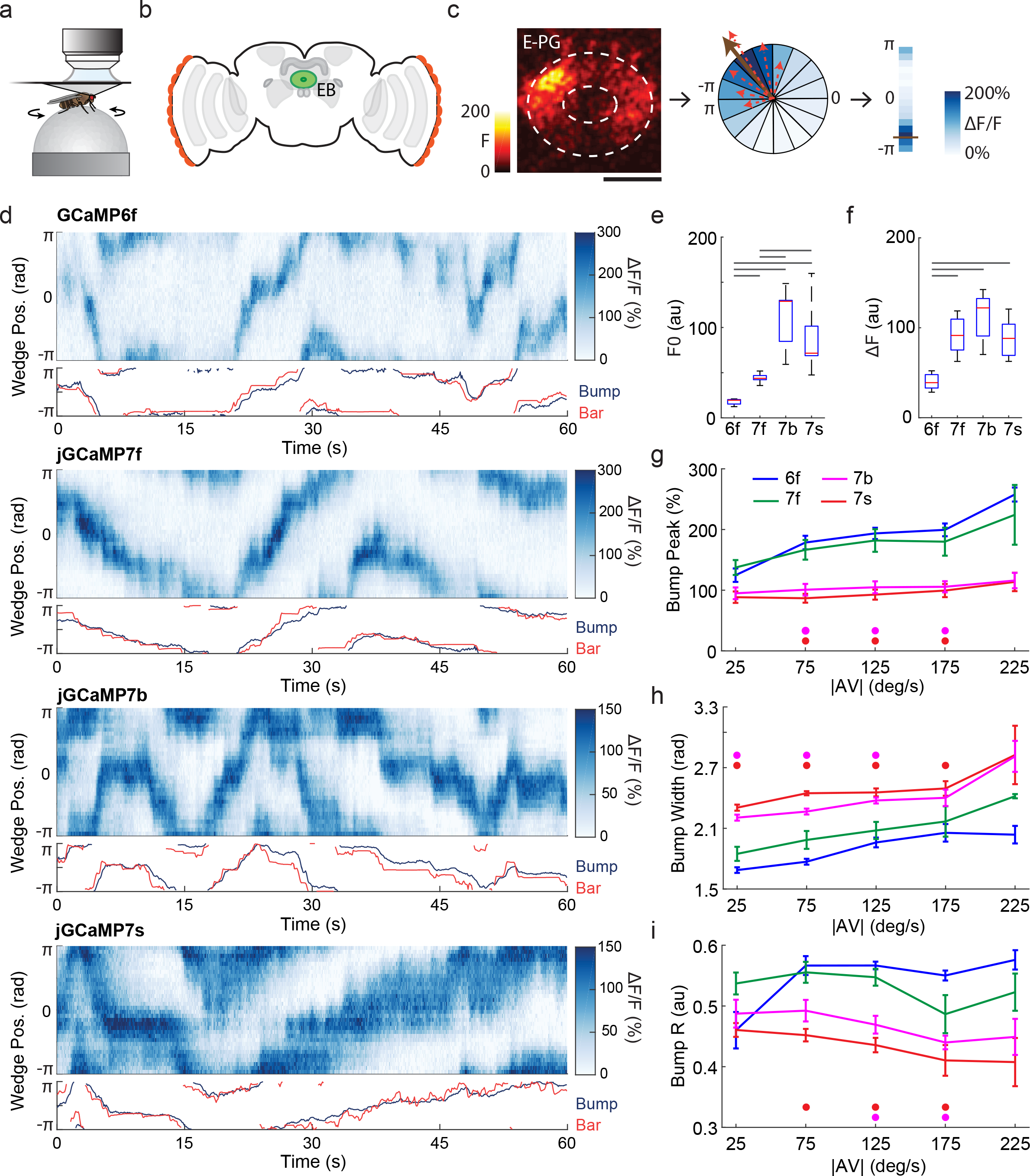
jGCaMP7 performance in adult *Drosophila* ellipsoid body neurons. **a**,A head-fixed fly walking on a spherical treadmill during two-photon calcium imaging. Flies walked in darkness or in closed-loop. During closed-loop trials, yaw rotations of the ball were used to update the azimuthal position of a 20-degree vertical bar presented on an LED display surrounding the fly. **b**,Schematic of the fly brain showing the ellipsoid body (EB, green). A population of neurons that encode the fly’s instantaneous heading direction (E-PG), innervate wedges of the EB, collectively tiling the whole structure. **c**,Imaging E-PG population activity in the EB. Z stacks were collected at 10 Hz, and maximum intensity projections were used to generate frames. Left, maximum intensity projection of EB calcium activity from a Z stack shows a single ‘bump’ of activity. Scale bar, 25 μm. Center, the EB was segmented into wedge-shaped ROIs (16 of 32 ROIs shown), and ΔF/F was computed for each ROI. ROIs are colored according to their ΔF/F (colorbar on right). For each frame, the population vector average (PVA; center, brown arrow) is computed by averaging the vectors from each ROI (for example, the dotted red arrows shown here for a few ROIs). The direction of the PVA is taken as the activity bump’s azimuthal position. **d**, Examples of E-PG calcium activity obtained using GCaMP6f, jGCaMP7f, jGCaMP7b, and jGCaMP7s (7 flies each; same data set used for all analyses in this figure). Top panels show E-PG calcium activity (blue) as a fly walks in closed loop with a 20-degree bar. The 32 EB ROIs are shown unwrapped from –π to π. Bottom panels show the azimuthal positions of the bump and 20-degree bar (offset removed, see methods). Notice that the amplitude and width of the bump, as well as the smearing that occurs during fast turns, all vary as a function of the sensor employed. **e**,Box plot showing the distribution of baseline fluorescence (F0) across flies for each sensor. The median across ROIs was taken as the fly’s F0. Horizontal bars mark pairwise comparisons with significantly different medians (p<0.01; Wilcoxon rank-sum tests). **f**,Same as in e, but for maximum ΔF. For each ROI, the difference between the 99^th^ and 1^st^ percentile was taken as the maximum ΔF. The median across ROIs was taken as the fly’s ΔF. **g**,Bump amplitude, (F_bump_ - F0)/F0*100, plotted as a function of absolute angular velocity. For each fly, frames containing a single bump were sorted into 50 deg/s bins and the median bump amplitude was computed. Displayed are the mean ± SEM across flies for each sensor. Colored dots mark sensors with significantly different bump peak compared to GCaMP6f at each angular velocity bin (p<0.01; Wilcoxon rank-sum tests). **h**, Same as in g, but for bump width, defined as full width at half maximum. **i**,Same as in g, but for the length of the normalized population vector average. Panels a-d were adapted from ^23^.

It is also critical that sensors have sufficient dynamic range and speed to report fast changes in neuronal activity. We compared the jGCaMP7 variants to GCaMP6f. Previous work has shown that the amplitude of the bump increases with the fly’s angular velocity^10,23,24^. Indeed, jGCaMP7f and GCaMP6f both reported monotonically increasing activity as a function of the fly’s angular velocity, whereas jGCaMP7b and jGCaMP7s showed a relatively reduced dynamic range (Fig. 4g). Next, as two proxies for sensor kinetics, we quantified how the width and strength (PVA, see methods) of the bump varied with the fly’s angular velocity. During fast turns, slow sensors cause the bump to ‘smear’, which increases its width and decreases its strength. In agreement with the culture (Fig. 2) and larval data (Fig. 3), jGCaMP7f and GCaMP6f had faster kinetics compared to jGCaMP7b and jGCaMP7s, as evidenced by their reduced bump with and increased bump strength across a range of angular velocities (Fig. 4h-i). Finally, as a measure of sensitivity we measured the width of the bump during slow turns (<50 deg/s; Fig. 4h, first bin). In agreement with previous data, jGCaMP7f and GCaMP6f were less sensitive than jGCaMP7b and jGCaMP7s, as shown by their decreased bump width during slow turns. Overall, because of its high baseline fluorescence, its high signal level, and its fast kinetics, jGCaMP7f appears to be the sensor of choice for imaging small, dim structures with fast kinetics. In addition, for applications where speed is less critical, jGCaMP7b and jGCaMP7s may be employed.

### Imaging neural populations in the mouse visual cortex

We tested the jGCaMP7 sensors in the mouse primary visual cortex (V1) *in vivo* and compared them to published GCaMP6 data^25^. The majority of V1 neurons can be driven to fire action potentials in response to visual stimuli in the form of drifting gratings^4,26,27^. V1 neurons were infected with adeno-associated virus (AAV) expressing one of the jGCaMP7 variants under the human *synapsin I* promoter (AAV-*Syn1*-jGCaMP7 variant) and imaged 16-45 days later using 2- photon microscopy through a cranial window. Cortical layer (L) 2/3 neurons showed green fluorescence in the neuronal cytoplasm. Gratings drifting in 1 of 8 different directions were presented as visual stimuli to the contralateral eye and fluorescence responses were recorded^3,4,19^. Visual stimulus-evoked fluorescence transients were observed for individual cells that were stable across trials and tuned to stimulus orientation (Fig. 5a-b). Orientation tuning was similar for all constructs tested (supp. Fig. 4). The dynamics of the sensory stimuli were tracked by sensor responses (Fig. 5b-c, Video 1-3). In agreement with the cultured neuron results, jGCaMP7f has faster kinetics than the other jGCaMP7 sensors and is comparable to GCaMP6f. jGCaMP7s has slower decay time than all the other sensors (Fig. 5d, Table 1, Methods).

**Figure 5.**
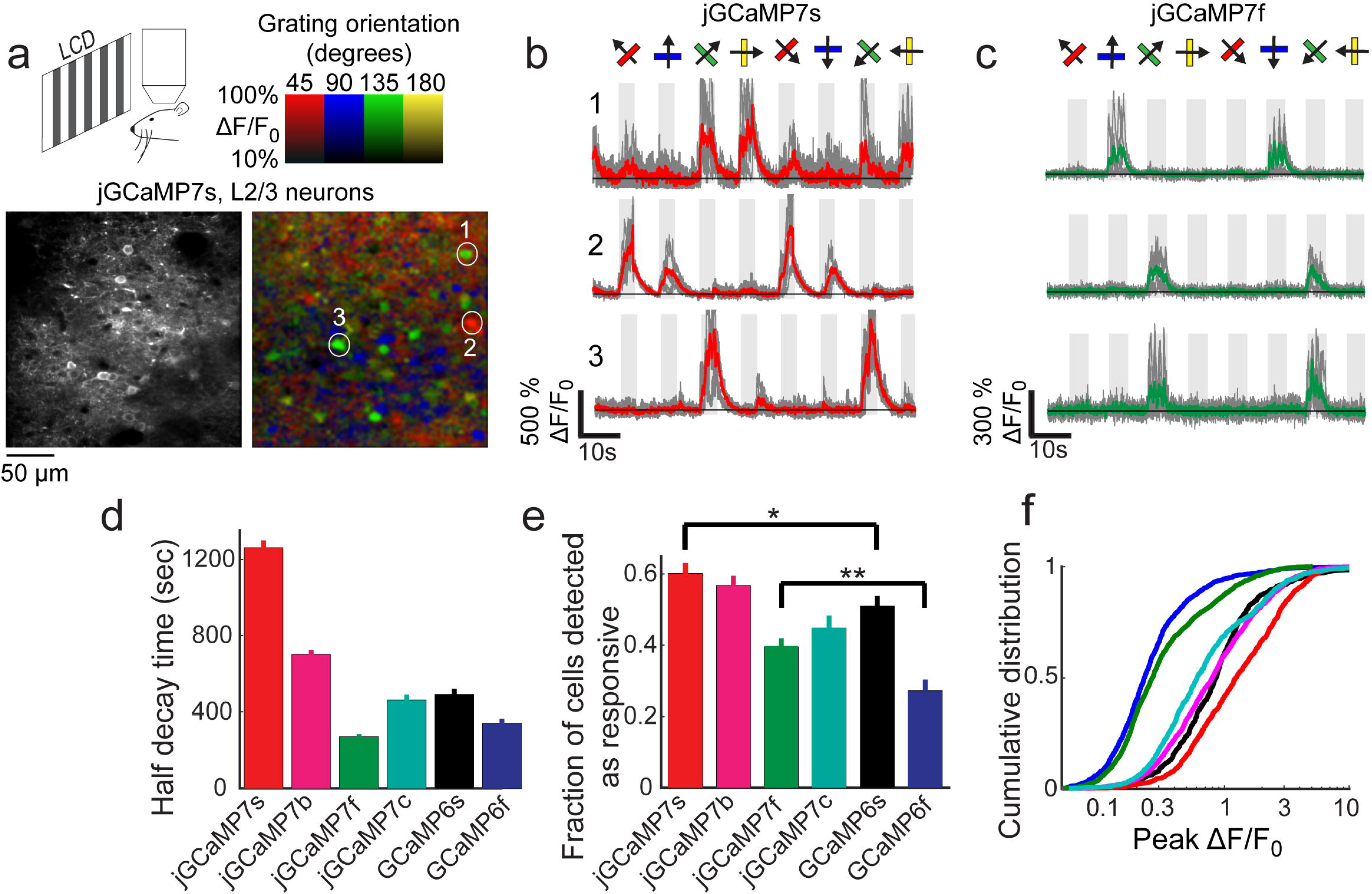
jGCaMP7 performance in the mouse primary visual cortex (V1) **a**,Top, schematic of the experiment. Bottom, image of V1 L2/3 cells expressing jGCaMP7s (left), and the same field of view color-coded according to the neurons’ preferred orientation (hue) and response amplitude (brightness). **b**,Example traces from three L2/3 neurons expressing jGCaMP7s. Single trials (gray) and averages of 5 trials (red) are overlaid. Eight grating motion directions are indicated by arrows and shown above traces. Each of the four orientations is color coded according to the color bar in panel **a**. The preferred stimulus is the direction evoking the largest response. jGCaMP7s traces correspond to the cells indicated in panel **a**(see also Video 1). Example traces from three L2/3 neurons expressing jGCaMP7f. Single trials (gray) and averages of 5 trials (green) are overlaid (see also Video 2). Half decay time of the fluorescence signal after the end of the visual stimulus for jGCaMP7s, 512 cells from n=3 mice; jGCaMP7f, 558 cells, n=5; jGCaMP7c, 282 cells, n=4; jGCaMP7b, 742 cells, n=3; GCaMP6s, 250 cells, n=3; GCaMP6f, 159 cells, n=3. **e**, Fraction of cells detected as responding to visual stimulus (ANOVA test, p<0.01) when expressing different calcium indicators. This fraction was significantly higher for jGCaMP7s vs. GCaMP6s and for jGCaMP7f vs. GCaMP6f respectively (Wilcoxon rank sum test; *, p<0.05; **, p<0.01). Error bars correspond to s.e.m (50 fields-of-view, jGCaMP7s; 55, jGCaMP7f; 40, jGCaMP7b; 38, jGCaMP7c; 23, GCaMP6s; 27, GCaMP6f) **f**, Distribution of ΔF/F amplitude for the preferred stimulus. the right-shifted curves of jGCaMP7s vs. GCaMP6s and jGCaMP7f vs. GCaMP6f, indicates significant enhancement of response amplitudes (p<0.001 Wilcoxon rank sum test. 75 percentile values of 250% and 54% vs. 130% and 36% for jGCaMP7s and jGCaMP7f vs. GCaMP6s and GCaMP6f). (980 cells, jGCaMP7s; 1623, jGCaMP7f; 1503, jGCaMP7b; 772, jGCaMP7c; 672, GCaMP6s; 873, GCaMP6f), same colors as in **f**.

We compared the sensitivity of the jGCaMP7 and GCaMP6 sensors^4,19^. One measure of sensitivity is the fraction of neurons detected as responsive to visual stimuli (Fig. 5e, ANOVA test, P < 0.01). For jGCaMP7s and jGCaMP7f, this fraction was significantly higher than for GCaMP6s and GCaMP6f (P=0.024 and 0.008, respectively, Wilcoxon rank-sum test). A second measure was the cumulative distribution of the peak ΔF/F_0_ amplitude from each cell (Fig. 5f), where a right-shifted distribution indicates enhanced sensitivity. This comparison also indicated that jGCaMP7s and jGCaMP7f are more sensitive than GCaMP6s and GCaMP6f (p<0.001, Wilcoxon rank-sum test). The 75th percentile ΔF/F_0_ amplitudes for jGCaMP7s and GCaMP6s were 250% and 130%, respectively, while for jGCaMP7f and GCaMP6f they were 54% and 36%, respectively. Finally, jGCaMP7b and jGCaMP7c demonstrated similar sensitivity to GCaMP6s but with the additional feature of brighter or dimmer baseline fluorescence (Fig. 2d).

### Imaging calcium signals in dendritic spines

We tested jGCaMP7b for imaging calcium signals in dendritic spines *in vivo* (Fig. 6). To trace and image dendrites of single cells, sparse expression of either GCaMP6s or jGCaMP7b was achieved in two ways. In one set of experiments we injected AAV expressing a Cre recombinase-dependent variant of the sensor together with a low-titer Cre-expressing AAV in mouse V1 (Fig. 6a, see methods)^4^. Animals were presented with full field drifting grating while dendrite imaging was performed (Fig. 6b-d). In each imaging session, a resonant mirror is directed in 3D to follow a few dendrites and the cell body acquiring large sections of contiguous dendrites (200-500um) at high speeds (~10-15Hz volume rate). We compared the number of spines that were detectable as an indication of baseline florescence (Fig 6e). As expected, jGCaMP7b had more traceable spines per micrometer of imaged dendrite (Fig. 6f; medians [CI]: 0.15 [0.13-0.19] vs. 0.26 [0.19-0.37] spines/μm for 6s and 7b respectively; n=sessions[animals]: 26[6] and 15[2] for 6s and 7b respectively, P=0.002, Wilcoxon rank-sum test). Second, we compared the number of tuned spines (ANOVA test, p < 0.01) as a correlate of indicator sensitivity. The number of tuned spines was also higher for jGCaMP7b (Fig. 6g; medians [CI]: 0.12 [0.07-0.16] vs. 0.16 [0.12-0.27] spines/μm for 6s and 7b respectively; n=sessions[animals]: 26[6] and 15[2] for 6s and 7b respectively, P=0.035, Wilcoxon rank-sum test).

**Figure 6:**
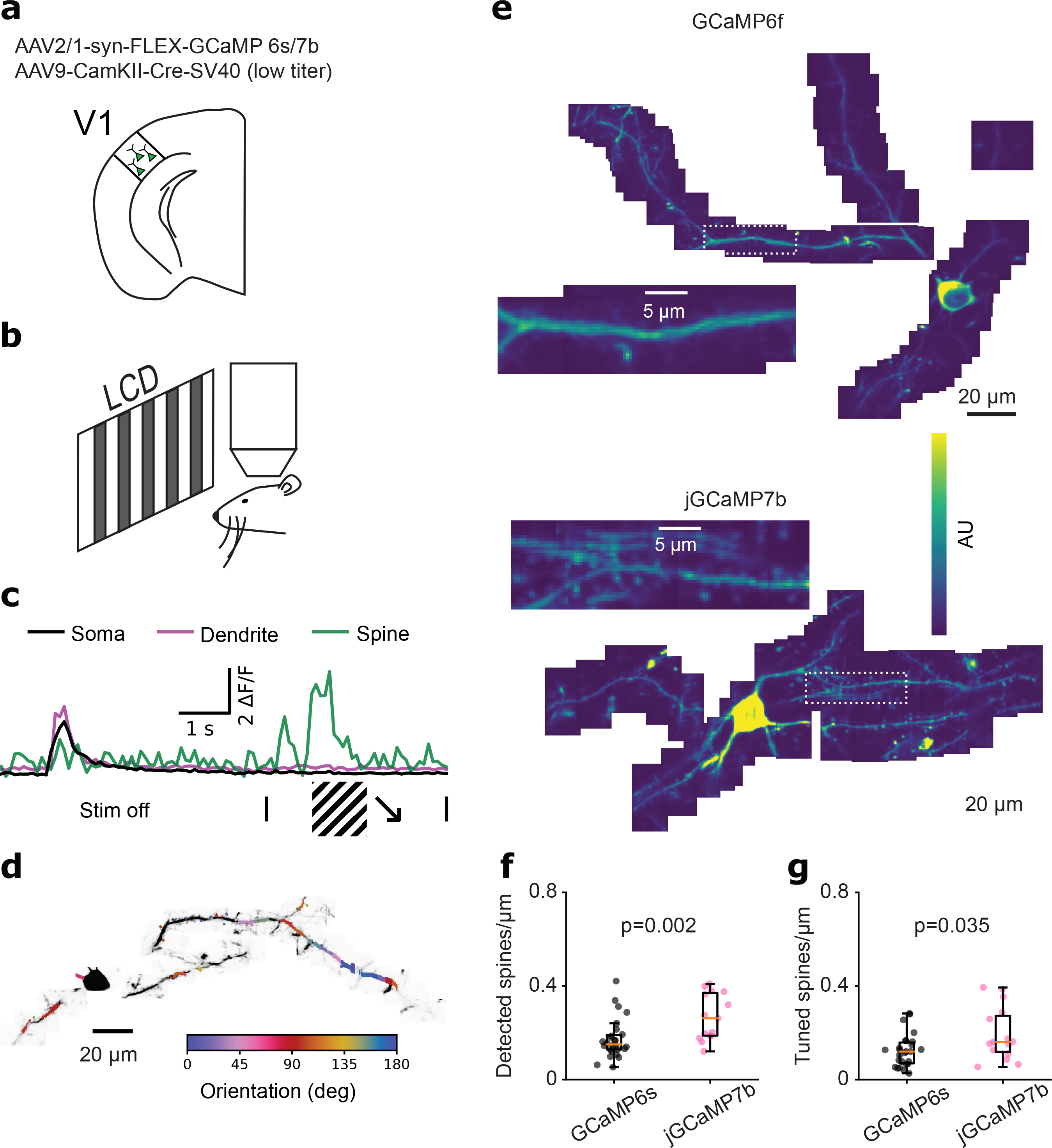
jGCaMP7b for improved dendritic spine imaging. **a.**Expression was achieved by injecting a mixture of low titer AAV expressing cre and high titer AAV expressing GCaMP in a cre-dependent manner. **b.**V1 imaging was performed while the mouse was passively viewing drifting gratings (Fig. 5a). **c.**Example traces from soma (black), dendrite (magenta) and a spine (green) before and during a presentation of drifting grating. **d.**Example imaging session Z-projected (gray) with the preferred orientation of each spine and dendritic segment overlaid in color. **e.**Z-projected mean images of example dendrite imaged in one session for GCaMP6s (top) and jGCaMP7b (bottom). Insets show the dotted white rectangles. **f.**jGCaMP7b allowed a higher density of detected spines (medians [CI]: 0.15 [0.13- 0.19] vs. 0.26 [0.19-0.37] spines/μm for 6s and 7b respectively; n=sessions[animals]: 26[6] and 15[2] for 6s and 7b respectively, P=0.002, Wilcoxon rank-sum test) **g.**jGCaMP7b also showed a higher density of spines tuned to orientation (medians [CI]: 0.12 [0.07-0.16] vs. 0.16 [0.12-0.27] spines/μm for 6s and 7b respectively; n=sessions[animals]: 26[6] and 15[2] for 6s and 7b respectively, P=0.035, Wilcoxon rank-sum test).

In a second set of experiments (Supp. Fig. 6), individual neurons in the upper cortical layers of mouse V1 were transduced with a plasmid expressing either GCaMP6s or jGCaMP7b using single-cell electroporation (Supp. Fig. 6a, Methods). As above, jGCaMP7b expressing neurons had higher densities of detectable spines (Supp. Fig 6d; median [CI]: 0.23 [0.17-0.32] vs. 0.3 [0.23-0.37] spines/μm for 6s and 7b respectively; n= 35 dendritic segments (5 neurons from 4 animals) and 37 dendritic segments (5 neurons from 4 animals) for 6s and 7b respectively, P=0.01, Wilcoxon rank-sum test).

The number of active spines was also higher for jGCaMP7b (Supp. Fig. 6e; medians [CI]: 0.17 [0.12-0.24] vs. 0.23 [0.21-0.33] spines/μm for 6s and 7b respectively; n= 35 dendritic segments (5 neurons from 4 animals) and 37 dendritic segments (5 neurons from 4 animals) for 6s and 7b respectively, P=0.001, Wilcoxon rank-sum test). The baseline brightness of jGCaMP7b allows for better detection of small neuronal structures and its high sensitivity facilitates detection of neural activity in these structures.

## Discussion

We developed new GCaMP calcium sensors with enhanced sensitivity, and with brightness and kinetic properties tailored to specific imaging applications. Under favorable conditions, GCaMP6 can detect single APs and can be imaged over multiple weeks^2,4^. However, when GCaMP6 is used under challenging conditions, for example at high imaging rates or low magnifications required for imaging large numbers of neurons, the fidelity for AP detection is reduced^5^. The enhanced SNR of jGCaMP7s and jGCaMP7f relative to GCaMP6s and GCaMP6f will permit *in vivo* imaging at higher speed and over wider fields of view. The greater brightness of jGCaMP7b enables more robust imaging in thin axonal and dendritic processes, where probe density, motion artifacts and photobleaching can be particularly challenging. The lower resting fluorescence of jGCaMP7c, while still possessing high SNR, will facilitate very large population imaging, as fluorescence from neuropil and inactive neurons will create less background to interfere with observation of active neurons. Imaging with jGCaMP7c will be helped by co-expression of a red fluorescent cellular marker to allow detection of jGCaMP7c-expressing neurons.

The improvement in detection of APs by the jGCaMP7 sensors was not accompanied by an increase in the maximal GCaMP fluorescence signal at saturating calcium. Tightening the indicator affinity for calcium caused saturation at lower numbers of spikes. Additional increases in affinity will further reduce the dynamic range. Future improvements might depend on increases in the GCaMP maximal fluorescence, which will require significant changes in the GCaMP scaffold, such as replacement of the cpGFP with brighter fluorescent proteins, redesign of the calmodulin-M13 interface with the cpGFP chromophore, or an entirely different fluorophore. Given the current GCaMP scaffold, we anticipate future improvements in indicator kinetics and indicators with red-shifted excitation spectra for 2-photon imaging with fiber lasers.

## Acknowledgements

We thank Aaron Kerlin and Dan Flickinger for design of the microscope used for spine imaging, Michael Reiser, Matthew Isaacson, Jim Chen, Jinyang Liu, and Andy Chiu for the G4.0 panel display system used in the fly-on-ball imaging experiments, and Deepika Walpita and Jenny Hagemeier for neuronal culture. This work is part of the GENIE project at Howard Hughes Medical Institute Janelia Research Campus. AK is supported by the Hertie Foundation.

## Online methods

All surgical and experimental procedures were in accordance with protocols approved by the HHMI Janelia Research Campus Institutional Animal Care and Use Committee and Institutional Biosafety Committee.

### Neuronal culture screen

The neuron-based screening system was previously described ^19,20^. In short, GCaMP variants were cloned into an expression vector contained a human synapsin I (SYN1) promoter for neuronal-specific expression, the tested variant, an internal ribosome entry site, a nuclearly-targeted mCherry, and a woodchuck hepatitis virus post-transcriptional regulatory element (WPRE). Neonatal rat hippocampi (P0) were dissociated, transfected with the expression vectors, and cultured in 96-well plates for 16-18 days. The plates were imaged using an EMCCD camera (Andor iXon DU897-BV, 35Hz) while trains of action potentials were triggered by field stimulation within each well. The acquired data were analyzed to segment somata and extract the fluorescence traces for single cells. Several parameters were extracted, such as the ΔF/F_0_ response amplitude, half-rise time, half-decay time, SNR, etc. For combining data from multiple wells for the same variant, the median values per well were averaged.

We conducted two cycles of screening experiments, and in the second screening cycle, the excitation intensities for GCaMP and mCherry fluorescence were increased to better detect low baseline fluorescence constructs. Illumination was provided by blue or white LEDs (GCaMP filter set, excitation: 450–490 nm, dichroic: 495 nm long-pass, emission: 500–550 nm; and mCherry filter set, excitation: 540–580 nm, dichroic: 585 nm long-pass, emission: 593– 668 nm). Illumination power was 2 mW and 7.5 mW for the GCaMP channel at the sample plane for the first and second screening cycles, respectively, and 3 mW for the mCherry channel. Results for the two screening cycles were merged to select best constructs. We merged parameters that were robust across the illumination change, including ΔF/F_0_ and kinetics. For each construct, we calculated the P-value (Wilcoxon rank sum test) for its improvement compared to GCaMP6s, under identical illumination conditions, and picked the higher value for the merged database. The kinetics of the selected jGCaMP7 sensors was measured in a third cycle with faster imaging rate (110Hz).

### Titrations and kinetic measurements of sensor proteins in solution

Sensors were purified, and calcium titrations, pH titrations, and stopped-flow measurements were performed essentially as described^4,19^. For protein purification, bacteria (T7 Express, NEB) were grown at 30°C for 48 h in 100 mL of Studier ZYM-5052 autoinduction media with 150 μg/mL ampicillin. Cells were resuspended in 4 mL of B-PER (ThermoFisher) and 16 mL of 20 mM Tris, pH 7.5, 300 mM NaCl, 1 mM imidazole. Cells were lysed by 90 s of sonication in 1 mg/mL lysozyme, and cell debris was removed by centrifugation. Lysate was run over 1000 μL of Ni^2+^-charged HisPur resin (Fisher) in gravity columns. Columns were first washed with 20 column volumes of 20 mM Tris, pH 7.5, 300 mM NaCl, 1 mM imidazole. They were then washed with 10 column volumes of 20 mM Tris, pH 7.5, 500 mM NaCl, 10 mM imidazole. GCaMP protein was eluted in 3000 μL of 20 mM Tris, pH 7.5, 100 mM NaCl, 100 mM imidazole.

For calcium titrations, GCaMP protein was first diluted 1:100 in triplicate in 30 mM MOPS, pH 7.2, 100 mM KCl with either 10 mM EGTA (zero free calcium) or 10 mM CaEGTA (~39 μM free calcium). These 2 solutions were mixed in various ratios to give 11 different free calcium concentrations (Calcium Calibration Buffer Kit #1, Life Technologies). GCaMP fluorescence (excitation 485 nm, 5 nm bandpass; emission 510 nm, 5 nm bandpass) was quantitated using a Safire^2^ plate reader (Tecan). The 0 and 39 μM free Ca^2+^ points were used for F_max_ and F_min_ determination. Calcium titration curves were fit to sigmoidal binding functions, and the Hill coefficient and K_d_ for Ca^2+^ for the GCaMP variant was extracted. Data were fit using Prism (GraphPad Software). Chromophore concentration was measured from the absorbance (447 nM) of protein denatured by 0.1 M NaOH (extinction coefficient 44,000 M^−1^cm^−1^).

For pH titrations, purified protein was diluted into pH buffers containing 20 mM citrate, 20 mM Tris, 20 mM glycine, 100 mM NaCl, and either 5 mM CaCl_2_ or 10 mM EGTA that were pre-adjusted to 36 different pH values between 4.0 and 12.0. The inflection point of a sigmoid fit to fluorescence versus pH was used to estimate pK_a_.

k_off_ and k_on_ were determined at room temperature using a stopped-flow device and fluorometer (Applied Photophysics). For k_off_, a sensor protein buffered in 1 μM free calcium was rapidly mixed with 30 mM BAPTA (both buffered in 30 mM MOPS, 100 mM KCl at pH 7.2). k_off_ values were determined from a single exponential fit to the signal decay, except for jGCaMP7s, which required a biexponential fit. For k_on_, a sensor protein (buffered in 30 mM MOPS, 100 mM KCl, 50 μM BAPTA at pH 7.2) was rapidly mixed (1:1) with free calcium at various concentrations produced through mixing of specific ratios of 2 mM BAPTA and 2mM calcium-BAPTA. The fluorescence change was measured during rapid mixing to obtain the observed rate constant (k_obs_) for calcium association for each free calcium concentration. The k_on_ value was determined by fitting the observed data to the equation k_on_ = (k_obs_-k_off_)/[Ca^2+^]^n^, where n was the Hill coefficient. Values are reported as mean ± s.e.m. where noted.

### Spectroscopy of purified proteins

#### Optical properties

Unless noted, all measurements were performed on purified protein in 30mM MOPS, 100mM KCl, pH 7.2 with either 10 mM CaEGTA (+Ca) or 10 mM EGTA (−Ca). Protein absorbance spectra were obtained in ±Ca on a UV-VIS spectrometer (Lambda 35, Perkin Elmer). These solutions were then denatured at pH 13 and the absorbance re-measured, where the protein chromophore concentration was determined using the extinction coefficient of denatured GFP (44,000 M^−1^cm^−1^ at 447nm). Extinction coefficients at pH 7.2 were then obtained from the measured absorbance at pH 7.2 and the measured chromophore concentration. Fluorescence emission and excitation spectra in ±Ca buffer (pH 7.2) were measured with a fluorimeter (LS 55, Perkin Elmer), from which the 1-photon fluorescence ΔF/F was determined. Quantum yield measurements were obtained using an integrating-sphere spectrometer (Quantaurus, Hamamatsu) for proteins in +Ca buffer. For proteins in −Ca buffer, quantum yield was determined by measuring relative fluorescence using GCaMP6f in −Ca buffer as reference (Φ = 0.66).

#### Two-Photon cross section

The two-photon excitation spectra were obtained as previously described^3,28^. Protein solutions in +Ca or −Ca buffer were prepared in a coverslip-bottomed dish (MatTek) and measured on an inverted microscope (IX81, Olympus) equipped with a 60X, 1.2NA water immersion objective (Olympus). The laser source was an 80 MHz Ti-Sapphire laser (Chameleon Ultra II, Coherent) scanned from 710 nm to 1080 nm, adjusted to deliver 1 mW average power at the sample across the excitation spectrum. At each excitation wavelength, fluorescence collected by the objective and passed through short-pass (720SP, Semrock) and bandpass filters (625BP90, Semrock), was detected in the image plane by a fiber-coupled avalanche photodiode (APD) (SPCM_AQRH-14, Perkin Elmer). Reference dye fluorescein at known concentration was prepared in pH 9.5 borate buffer and measured in the same experimental run. Action cross-sections (AXS), the product of quantum yield and 2-photon cross section, were calculated at each wavelength for the GCaMP indicators using the equation

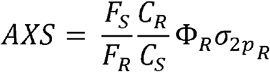

where subscripts S and R refer to GCaMP sample and dye reference, respectively, *F* is the measured fluorescence rate, *C* is concentration, Φ_R_ and 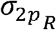 are the fluorescence quantum yield and 2-photon cross section of the reference dye. For fluorescein we use cross section data from^29^, and separately from^30^, and average these at each wavelength to obtain the GCaMP 2- photon action spectrum.

#### Two-photon molecular brightness

Fluorescence correlation spectroscopy (FCS) was used to determine the 2-photon molecular brightness (fluorescence rate per molecule) of the GCaMP indicators^28^. In FCS, samples are illuminated and the fluorescence count rate detected by the APD is fed into an autocorrelator (Flex03LQ, Correlator.com). The fluorescence autocorrelation obtained is fit to a simple diffusion model written and executed in Matlab^28^, in order to calculate the average number of molecules < *N* > present in the focal volume. The 2-photon molecular brightness *ϵ* (kilocounts/sec/molecule, or kcpsm) at a specific laser power is determined by the average fluorescence rate <*F*> divided by the number of molecules <*N*>, *ϵ* = 〈*F*〉/〈*N*〉. As a function of laser power, the molecular brightness initially increases, peaks, then decreases at high power. The peak molecular brightness *ϵ*_*max*_ is determined from a plot of E versus laser power, and was used to characterize the relative photostability of the GCaMP indicators. To determine peak brightness, protein solutions at 50-100 nM were prepared in +Ca buffer and excited at 930 nm, and an FCS run performed (200 sec duration) at each of several laser powers from 2–30 mW.

### Imaging of the *Drosophila* larval neuromuscular junction

We made w^1118^; ;PBac[20XUAS-IVS-Syn21-op1-GECI-p10]VK00005 transgenic flies and crossed them with w^1118^; ;R57C10-Gal4 in VK00020, R57C10-Gal4 in VK00040 pan-neuronal driver line. Sensor cDNAs were individually codon-optimized for Drosophila for improving expression^31^.

The NMJ assay is similar to the one used previously^4,19^. Briefly, female actively crawling 3rd instar larvae of the progenies were dissected under minimum illumination intensity, and type Ib boutons on muscle 13 from segment A3-A5 were wide-field imaged in HL-6 saline while corresponding axons were electrically stimulated with a suction electrode driven by a customized stimulator. The temperature and pH were monitored simultaneously with the stimulation and imaging. The HL-6 saline contains 2mM of calcium. In addition, 7mM of L-glutamic acid was added to prevent muscle contraction by saturating the glutamatergic synapse of NMJ. A mercury lamp (X-CITE exacte) light source was used for excitation and out-of-objective power of less than 5mW was used. The light intensity was calibrated so that no significant photo-bleaching was detected during each trial. The filters for imaging were: exciter: 472/30; dichroic: 495; emitter: 520/35. EMCCD cooled to −70 C was acquiring at 30 fps. Data were analyzed in MATLAB (MathWorks).

### Two-photon imaging of E-PG population activity in head-fixed, walking adult *Drosophila*

We crossed the w^1118^; ;PBac[20XUAS-IVS-Syn21-op1-GECI-p10]VK00005 transgenic flies described in the NMJ assay to a split-Gal4 driver, SS00096, that selectively expresses in E-PG neurons ^32^. Flies (females, age 6-9 days, n=7 flies per group) were prepared for imaging as previously described^10,33^. Briefly, flies were anaesthetized at 4-degrees, their proboscis immobilized with wax to reduce brain movements, and their head/thorax UV-glued to a holder with a recording chamber. To gain optical access to the brain, a section of cuticle between the ocelli and antennae was removed, along with the underlying fat and air sacs. Muscle 16 was cut to reduce pulsatile brain movements. Throughout the experiment, the head was submerged in saline containing (in mM): NaCl (103), KCl (3), TES (5), trehalose (8), glucose (10), NaHCO_3_ (26), NaH_2_PO_4_ (1), CaCl_2_ (2.5), and MgCl_2_ (4), with a pH of 7.3 and an osmolarity of 280 mOsm.

Calcium imaging was performed with a custom-built two photon microscope controlled with ScanImage 2017 (Vidrio Technologies^34^). Excitation of GECIs was generated with an infrared (930 nm), femtosecond-pulsed (pulse width ~ 110 fs) laser (Chameleon Ultra II, Coherent) with 15 mW of power, as measured after the objective (20X Olympus XLUMPLFLN, 1.0 NA, 2.0 mm WD). Fast Z stacks (5 planes with 6 μm spacing and 3 fly-back frames)were collected at 10 Hz by raster scanning (128×128 pixels; ~60×60 μm) using an 8 kHz resonant-galvo system and piezo-controlled Z positioning. Focal planes were selected to cover the full extent of E-PG processes in the EB. Emitted light was directed (primary dichroic: 735; secondary dichroic: 594), filtered (filter A: 680 SP; filter B: 514/44), and detected with a *GaAsP* PMT (H10770PB-40, Hamamatsu).

Following dissection, flies were positioned on an air-supported polyurethane foam ball (8 mm diameter, 47 mg) under the microscope and allowed to walk. Rotations of the ball were tracked at 500 Hz, as described previously^33^. Behavioral data and imaging timestamps were recorded using WaveSurfer (http://wavesurfer.janelia.org/). Side and rear views of the fly were recorded from two CMOS cameras (Grasshopper3, FLIR) using BIAS (IO Rodeo). For each fly, we concatenated the data from two to eight, one-minute trials for darkness and closed-loop periods separately. During closed-loop trials, yaw rotations of the ball were used to update the azimuthal position of a bright, 20-degree vertical bar presented on an LED display surrounding the fly. All experiments used unity gain, where one full yaw rotation of the ball produced a 360-degree rotation of the bar. The display system was constructed from an updated version (referred to as G4.0) of the LED panels previously described ^35^. The visual arena was composed of 12 panels, similar to those previously describes, but designed for higher display speeds and higher pixel density. The visual arena covered 240 degrees in azimuth and ~60 degrees in elevation using a grid of 192 × 64 pixels (green LEDs; emission peak: 565◼nm; 2.5 mm pixel spacing), and was refreshed synchronously at 1000 Hz. The diameter of each pixel’s subtended area is at most 1.25 degrees on the fly eye.

Data analysis was performed with MATLAB (MathWorks) and the CircStat toolbox ^36^. Each Z-stack was reduced to a single frame using a maximum intensity projection (**Fig. 4c**, left). An ellipse was manually drawn around the perimeter of the ellipsoid body (EB) and automatically segmented into 32 equal-area, wedge-shaped regions of interest (ROIs; **Fig. 4c**, right). The number of ROIs was chosen to match the number of anatomically-define, EB demi-wedges ^22^. Activity within each ROI was averaged for each frame, producing 32 ROI time series (**Fig. 4d**). For each ROI time series, baseline fluorescence (F0) was define as the average of the lowest 10% of samples (**Fig. 4e**). ΔF/F was computed as (F-F0)/F0*100, where F is the instantaneous fluorescence from the raw ROI time series. To quantify each ROI’s maximum ΔF, we used the difference between the 99^th^ and 1^st^ percentiles (**Fig. 4f**). To quantify the bump’s peak ΔF/F, width, and strength (**Fig.s 4g-i**), we first detected whether a single bump of activity was present on a frame-by-frame basis, as previously described ^10^. Briefly, each ROI’s ΔF/F time series was filtered with a 3rd order Savitzky–Golay filter over 7 frames. Then, for each frame, a single bump of activity was present if there existed a single, contiguous set of ROIs whose ΔF/F was greater than 1 SD above the mean across all ROIs. For each frame with a single bump of activity, the ROI with the largest ΔF/F was used to determine the bump ΔF/F (**Fig. 4g**). Bump width (**Fig. 4h**) was defined as the full width at half max. Finally, we used the population vector average (PVA) as a measure of bump strength (**Fig. 4i**). For each frame with a single bump, PVA was computed by taking the circular mean of vectors whose angles were the ROIs’ wedge positions and whose length was equal to the ROIs’ ΔF/F. The magnitude of this mean resultant vector length was normalized to have a maximum possible length of 1. Each fly’s bump amplitude, bump width, and normalized PVA length was taken as the average across frames and plotted as the mean ± SEM across flies (**Fig.s 4g-i**). As described previously ^10^, there is a fly-specific angular offset between the position of the bump in the EB and the position of the bar stimulus. This difference was subtracted in the bottom panels of **Fig. 4d**for display purposes.

### *In vivo* imaging of mouse visual cortical neurons

*In vivo* imaging and data analysis of mouse V1 neurons were performed as described in^4,19^. Mice were transcranially injected with AAV-SYN1-jGCaMP7 construct. Mice were anesthetized using isoflurane (2.5% for induction, 1.5% during surgery). A circular craniotomy (2–2.5 mm diameter) was made above V1 (centered 2.7 mm left, and 0.2 mm anterior to Lambda suture) and covered with thin 1% agarose layer. A 3 mm diameter round glass coverslip (no. 1 thickness, Warner Instruments) was cemented to the craniotomy using black dental cement (Contemporary Ortho-Jet). A custom titanium head post was cemented to the skull. The mice were taken to the imaging setup immediately after the completion of the cranial window implantation. They were kept on a warm blanket (37°C) and anesthetized using 0.5% isoflurane and sedated with chlorprothixene (20–30 l at 0.33 mg/ml, i.m.).

Imaging was performed with a custom-built two-photon microscope with a resonant scanner. The light source was an Insight femtosecond-pulse laser (Spectra-Physics) running at 940 nm. The objective was a 16x water immersion lens with 0.8 NA (Nikon). Images were acquired using ScanImage 5 (vidriotechnologies. com)^37^. Functional images (512×512 pixels, 250×250 mm^2^) of L2/3 cells (100–250 mm under the pia) were collected at 29 Hz. Laser power was up to 50 mW at the front aperture of the objective.

Visual stimuli were moving gratings generated using the Psychophysics Toolbox^38,39^ in MATLAB (Mathworks), presented using an LCD monitor (30×40 cm), placed 25 cm in front of the center of the right eye of the mouse. Each stimulus trial consisted of a 4 s blank period (uniform gray display at mean luminance) followed by a 4 s drifting sinusoidal grating (0.05 cycles/degree, 1 Hz temporal frequency, 8 different directions). The stimuli were synchronized to individual image frames using frame-start pulses provided by ScanImage 5. The monitor subtended an angle of ±38° horizontally and ±31° vertically around the eye of the mouse.

The acquired data was analyzed using MATLAB. Regions of interest (ROIs) corresponding identifiable cell bodies were selected using a semi-automated algorithm, and the fluorescence time course was measured by averaging all pixels within the ROI, after correction for neuropil contamination (r=0.7), as described in detail in^4^. For correct neuropil subtraction, we excluded from the analysis cells with somatic signal lower than 103% of the surrounding neuropil signal. We used ANOVA test (P=0.01) for identifying cells with significant increase in their fluorescence signal during the stimulus presentation (responsive cells). We calculated ΔF/F_0_=(F-F_0_)/F_0_, where F is the instantaneous fluorescence signal, and F_0_ is the average fluorescence 0.7 sec before the start of the visual stimulus. For each responsive cell we defined the preferred stimulus as the stimulus which evoked the maximal ΔF/F_0_ amplitude (averaging the top 25% of ΔF/F_0_ values during the 4sec of stimulus presentation). The half decay time (Fig. 5d, Table 1) was calculated as following, for each responsive cell we averaged its ΔF/F_0_ response to the preferred stimulus over 5 trials. We also calculated the standard deviation of the averaged baseline signal during 1 sec before the start of the stimulus. Only cells with maximal ΔF/F_0_ amplitude which was higher than 4 standard deviations above the baseline signal were included in the analysis (512 cells for jGCaMP7s; 558, jGCaMP7f; 742, jGCaMP7b; 282, jGCaMP7c; 250, GCaMP6s; 159, GCaMP6f). The time required for each trace to reach half of its peak value (baseline fluorescence subtracted) was calculated by linear interpolation. The fraction of cells detected as responsive (Fig. 5e) was calculated as the number of significantly responsive cells over all the cells that were analyzed. The cumulative distribution of peak ΔF/F_0_ responses (Fig. 5f) included the maximal response amplitude from all analyzed cells, calculated as described above for each cell’s preferred stimulus (n=980 cells from n=4 mice for jGCaMP7s; 1623, n=5, jGCaMP7f; 1503, n=3, jGCaMP7b; 772, n=4, jGCaMP7c;).

### In-vivo imaging of dendrites and spines in mouse V1

A circular (∼2.5 mm diameter) craniotomy was made above left V1 (centered at 0.5 mm anterior and 2.7 mm lateral to lambda). Animals were injected a high titer (~1e11) CRE-dependent calcium indictor (AAV2/1-syn-FLEX-GCaMP6s, AV-1-PV2819, n=6 animals; AAV2/1-syn-FLEX-GCaMP7b, AddGene #104493, n=2 animals) combined 1:1 with low titer (~1e7) CRE (AAV9-CamKII-Cre-SV40) to achieve sparse expression. Stimulus was generated using psychopy (Peirce, 2009) on a calibrated monitor at ~15cm form the right eye of the animal and at an angle of ~30 degrees from the long axis of the animal. The screen occupied around 120×80 (horizontal × vertical) degrees of the animal’s visual field. Stimulus consisted of full screen square wave drifting gratings (0.03 cycles/degree, 100% contrast, 2Hz) at 8 directions for 4s. Between stimuli, 6s of a gray screen was presented to allow the slow decay of the indicator to return to baseline. The stimulus was randomized and repeated for 12-30 times in each session.

A custom high NA resonant-galvo-galvo microscope was used to preform morphological and 3D targeted dendritic imaging. Briefly, a morphological stack was taken around an injection site and cell bodies were manually traced using neuromantic^40^. Cells were selected based on a reliable retinotopic response and sufficient baseline fluorescence to enable tracing of dendritic branches. Selected cells were manually traced using neuromantic and each day a subset of dendrites was selected. The cell’s reconstruction was loaded to a custom Matlab GUI (Mathworks, Natick, MA) and was imagined using ScanImage 2017 (Vidrio Technologies, Ashburn, VA). All frames in the volume were 72×36 pixels/lines and spanned 24×12um giving 3px/um resolution and taking ~2ms to scan (@16kHz line rate). The number of dendritic segments was selected to attain a volume rate of ~10Hz. Motion correction and time course extraction procedures were used. Briefly, a cross-correlation based approach was combined with an iterative based optimization to correct for x-y motion and estimate the effect of z-motion on baseline fluorescence. After registration, an aligned averaged 3d stack was generated for manually tracing the dendritic centerline and the spines in each session using a custom Matlab GUI. Baseline was estimated by using a sliding window of 2000-time points (~2min) and fitting a Gaussian model to the F fluctuations of each ROI. This also considered the average target and number of expected pixels given the x, y, z shift assigned to each time point. Finally:

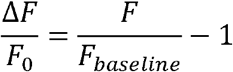

Noise was estimated for each ROI and time point by considering a Poisson process and estimating the number of pixels and average value of a single photon in our acquisition hardware. Significantly tuned spines were spines that had a p<0.01 for Kruskal–Wallis one-way ANOVA on ranks for the time averaged responses to all orientations. Mean z-projected images (Fig. 6a) were taken from time points where the soma’s DF/F values were lower than twice the estimated noise. This was done to avoid conflicting activity and baseline florescence.

### Targeted single-cell electroporation and two-photon imaging in mouse V1

The procedure for targeted single-cell electroporation of GCaMP6s (pGP-CMV-GCaMP6s) and jGCaMP7b (pGP-CMV-jGCaMP7b) plasmids was a modified version of a previously described method^41^. The experiments were performed in 8 C57BL/6 mice (4 for each plasmid) that were 45 to 60 days old. First, a head holder was implanted on the skull of each mouse under isoflurane anesthesia (2% for induction, 1.5% for surgery) and after a treatment with analgesics (xylocaine 2%; metamizole, 200 mg/kg; metacam, 1.5 mg/kg). Three days later, at the day of electroporation, dexamethasone (2mg/kg) was injected 2 hours before surgery. A square shaped craniotomy (1.5 mm by 1.5mm) was dissected over V1. For plasmid electroporation, a standard patch pipette with a resistance of 5-6 MΩ was used. The pipette was filled with an intracellular solution (consisting of 135 mM K-gluconate, 4 mM KCl, 10 mM HEPES, 4 mM Mg-ATP, 0.3 mM Na_2_-GTP, 10 mM Na-Phosphocreatine) containing ~0.1 pmol/μl of one of the plasmids. The intracellular solution contained in addition 100 μM OGB-1 for visualizing and navigating the pipette in the brain. The targeted single-cell electroporation procedure was performed in two-photon microscope under visual control in vivo ‘shadow-patching’^41^. Once the tip of the pipette was in contact with the cell membrane, a train of voltage pulses ^41^ was given to deliver the plasmid. The craniotomy was sealed by Kwik-Sil and Kwik-cast after electroporation. The actual imaging experiments were performed 48 hours after electroporation. For this purpose, the dura mater was removed and the brain was covered with agarose (2%). Dendrites and the cell body were imaged at 100 Hz frame rate at 920 nm two-photon excitation. Data processing was performed in MATLAB and included down sampling and movement correction. A baseline image was generated for each recording by averaging frames from periods without dendritic activity. Dendritic segments were traced, and ROIs were made on spines manually on the baseline image with a custom GUI. The fluorescence signals from the ROIs were extracted and spontaneous calcium transients (df/f) were collected through an automatic detection procedure.

### Reagent distribution

DNA constructs and AAV particles jGCaMP7 variants were deposited for distribution at Addgene (http://www.addgene.org). *Drosophila* stocks were deposited at the Bloomington Drosophila Stock Center (http://flystocks.bio.indiana.edu).

